# TECPR2 maintains mitochondrial homeostasis in neurodegeneration

**DOI:** 10.1101/2025.09.04.674193

**Authors:** Madhuri Chaurasia, Milana Fraiberg, Nemanja Subic, Oren Shatz, Kamilya Kokabi, Olee Gogoi, Olena Trofimyuk, Bat Chen Tamim-Yecheskel, Saskia Freud, Alik Demishtein, Ekaterina Kopitman, Inna Goliand, Sabita Chourasia, Yoav Peleg, Elena Ainbinder, Nili Dezorella, Zvulun Elazar

## Abstract

HSAN9 is a rare progressive neurodegenerative disease in children linked to bi-allelic loss-of-function mutations in the *TECPR2* gene. TECPR2 is a multi-domain protein harboring N-terminal WD repeats and C-terminal TECPR repeats, followed by a functional LIR motif that serves in autolysosomal targeting. Here, we show that the lack of TECPR2 leads to impairment of mitophagy that can be recovered by the expression of its C-terminal domain. Accordingly, we uncover severe mitochondrial dysfunction and accumulation of mitochondrial content in primary fibroblasts derived from an HSAN9 patient, and in embryonic fibroblasts and dorsal root ganglia derived from an HSAN9 mouse model. Strikingly, these mitochondrial defects are mediated by a mitochondrial stress through activation of the integrated stress response (ISR), whereas mitochondrial function is recovered by pharmaceutical or genetic suppression of ISR. Our findings provide a new link between mitophagy and ISR in mitochondrial homeostasis during neurodegeneration.

## Introduction

Hereditary sensory and autonomic neuropathy IX (HSAN9), previously known as Spastic paraplegia type 49 (SPG49; OMIM 615031), is a unique recessively inherited neuronal disorder linked to several mutations on TECPR2^1–3^. HSAN9 patients have intellectual disability, evolving spasticity, sensory-autonomic neuropathy, and dysmorphic appearance^1,4^.

Tectonin beta-propeller repeat-containing 2 (TECPR2) is a multi-domain protein with multiple WD repeats on its N-terminus, and six TECPR repeats followed by a functional LC3-interacting region (LIR) on its C-terminus^1,5^. TECPR2 is implicated in autophagy due to its interaction with ATG8 family proteins^6^, biogenesis of lysosome-related organelles complex 1 (BLOC1) and homotypic fusion and protein sorting (HOPS) protein complexes^5^, and the soluble N-ethylmaleimide-sensitive factor-attachment protein receptors (SNARE) protein vesicle-associated membrane protein 8 (VAMP8) that mediate autophagosome-lysosome tethering^7^.

Mitophagy, a selective elimination of damaged mitochondrial fragments by autophagy, plays a vital role in cellular homeostasis and is often affected in neurodegeneration^8,9^. Mitophagy is activated upon mitochondrial stress, including mitochondrial unfolded protein response (UPR^mt^), which results in mitochondrial biogenesis and mitophagy, when both processes are activated by phosphorylation of the α subunit of eukaryotic initiation factor 2 (eIF2α) through HRI kinase^10,11^. This phosphorylation leads to translational and transcriptional signaling induction by activation of stress-responsive transcription factors, such as activation transcription factor 4 (ATF4) and activation transcription factor 5 (ATF5)^12^.

Here, we show that lacking TECPR2 leads to aberrant mitochondrial homeostasis, elevated integrated stress response (ISR), and induced mitochondrial biogenesis. The reintroduction of TECPR2 successfully recovers both mitochondrial homeostasis and mitophagy. Moreover, we find that TECPR2 is required for mitochondrial clearance by parkin-dependent mitophagy, while other selective autophagy processes, such as ER-phagy and pexophagy, remain unaffected. Strikingly, mitochondrial homeostasis (but not mitophagy) is recovered by inhibition of ISR, ascribing a role for the latter in mitochondrial collapse in neurodegeneration.

## Results

### TECPR2 maintains mitochondrial homeostasis

Mitochondrial dysfunction is implicated in several neurodegenerative diseases, including PD, AD, ALS, and HD^13–15^. To test the mitochondrial state in HSAN9, we employed human primary fibroblasts derived from HSAN9 patients carrying heterozygous mutations on *TECPR2* alleles (Ex. 8+16), and mouse embryonic fibroblasts (MEF) cells derived from the HSAN9 mouse model^16^. Mitochondrial respiration of primary fibroblast cells and MEFs was determined using the Seahorse mitostress assay^17^. We observed reduced oxidative phosphorylation and low ATP production in the primary fibroblasts derived from HSAN9 patients and in MEF cells derived from TECPR2 knockout (KO) mice, indicating a severe mitochondrial dysfunction (Fig.1A and Fig. S1A). This was accompanied by the accumulation of mitochondrial reactive oxygen species (mtROS) visualized through MitoSOX Red by confocal microscopy (Fig. 1B) or fluorescently measured by flow cytometry (Fig. S1B and Fig. S1C). Treatment of wild-type (WT) and *TECPR2* KO MEF cells with the oxidative phosphorylation uncoupler carbonyl cyanide m-chlorophenylhydrazone (CCCP) resulted in mtROS accumulation only in the WT cells, whereas their level in the *TECPR2* KO cells was already saturated (Fig. S1D).

**Figure 1.**
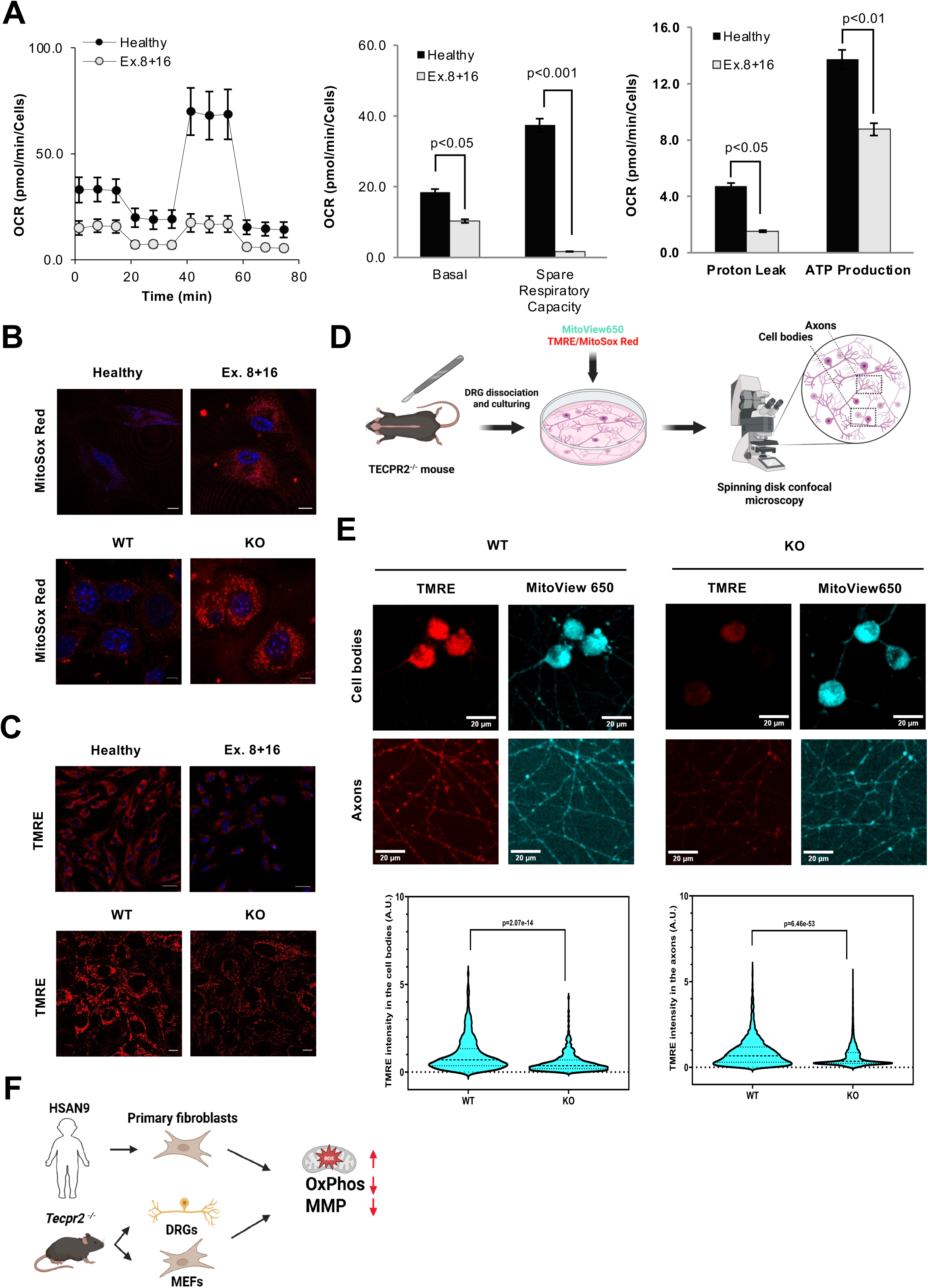
TECPR2 maintains mitochondrial homeostasis. **A.** Seahorse analysis for quantification of mitochondrial respiration in HSAN9 patient-derived fibroblasts versus healthy fibroblast cells. **B.** Confocal microscopy visualization of mtROS using MitoSox Red staining in HSAN9 patient-derived fibroblasts versus healthy control and MEF cells derived from WT and TECPR2 KO mice. Scale bar: 20 μm. **C.** Confocal microscopy visualization of mitochondrial membrane potential by TMRE staining in HSAN9 patient-derived fibroblasts, healthy controls, and MEF cells derived from WT and TECPR2 KO mice. Scale bar: 100 μm for human fibroblasts and 20 μm for MEF cells. **D.** Schematic presentation of DRGs culturing and imaging. **E.** Confocal microscopy visualization of mitochondrial membrane potential by TMRE staining in cell bodies and axons of DRGs derived from WT and TECPR2 KO mice. Scale bar: 20 μm. Statistical analysis was conducted using the Kruskal–Wallis test followed by Dunn’s post-hoc test; *\*\*\*\**p < 0.0001, ns: insignificant. **F.** Schematic illustration of disrupted mitochondrial homeostasis in TECPR2 KO cells.

To estimate the mitochondrial membrane potential (ΔѰm), human primary fibroblasts and MEFs were treated with tetramethylrhodamine ethyl ester (TMRE). Analysis by confocal microscopy (Fig. 1C) and flow cytometry (Fig. S1E and Fig. S1F) indicates mitochondrial depolarization in TECPR2 KO cells. Importantly, treating *TECPR2* KO MEFs with the mtROS scavenger MitoTEMPO led to significant recovery of mitochondrial membrane potential in TECPR KO cells, without affecting that of WT cells (Fig. S1G).

Since HSAN9 primarily affects neurons, we next investigated whether mitochondrial defects also exist in the neuronal system. Dorsal root ganglia (DRG) obtained from WT and TECPR2 KO mice were co-stained with TMRE and MitoView 650 to assess mitochondrial membrane potential and morphology. Spinning disk confocal microscopy was used to analyze both cell bodies and axons, as depicted in Fig. 1D. Here, too, we observed mitochondrial depolarization in both cell bodies and axons of TECPR2 KO DRGs (Fig. 1E).

These findings indicate that depletion of TECPR2 leads to severe mitochondrial dysfunction, including reduced OxPhos, accumulated mtROS, and mitochondrial depolarization (Fig. 1G). TECPR2, therefore, plays an essential role in mitochondrial homeostasis.

### TECPR2 supports mitophagic flux upon mitochondrial stress

It has been established that a reduction in mitochondrial membrane potential leads to activation of mitophagy^18^. To determine whether mitophagy was affected upon depletion of TECPR2, mitochondria-targeted mKeima (mt-mKeima), a reporter of autophagic flux^19^, was stably introduced into MEF cells obtained from WT or TECPR2 KO mice. The cells were treated with the mitochondrial uncoupler CCCP to induce mitophagy and analyzed by confocal microscopy^20^ and flow cytometry^21^. As indicated in Fig. 2A and Fig. S2A, treating cells with CCCP led to activation of mitophagy in WT but not in TECPR2 KO cells.

**Figure 2.**
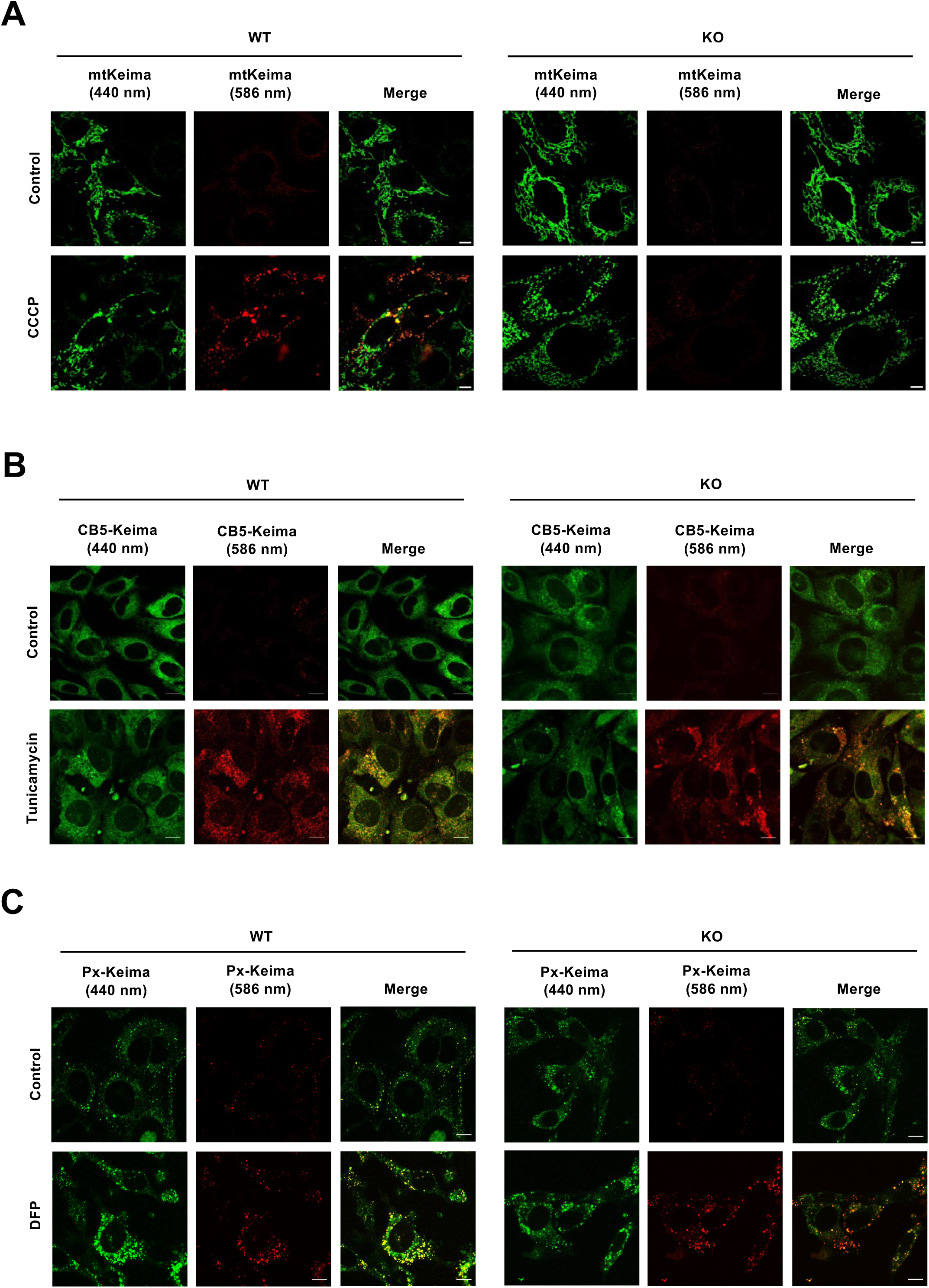
TECPR2 supports mitophagic flux upon mitochondrial stress. **A.** Confocal microscopy visualization of mitophagy in live TECPR2 KO and WT cells stably expressing mKeima under basal and mitophagy-inducing conditions by CCCP for 4 h. **B.** Confocal microscopy visualization of ERphagy in live TECPR2 KO and WT cells expressing Cb5-Keima under basal and ERphagy-inducing conditions by 1µM tunicamycin for 4 h. **C.** Confocal microscopy visualization of pexophagy in live TECPR2 KO and WT cells stably expressing Px-Keima under basal and pexophagy-inducing conditions by 1mM DFP treatment for 24 h.

Given the potent mitophagic inhibition, we next set out to determine whether other autophagy pathways, such as ERphagy and pexophagy, are affected by the loss of TECPR2. To this end, WT and KO MEFs stably expressing mKeima targeted by cytochrome B5 to the ER lumen (Cb5-mKeima) or targeted by SKL to peroxisomes were established (Px-mKeima). As depicted in Fig. 2B and Fig. S2B, ERphagy induced by tunicamycin remained unaffected upon depletion of TECPR2. Similar results were observed when pexophagy was tested in cells stably expressing Px-mKeima and treated with Deferiprone (DFP) (Fig. 2C and Fig. S2C). These data support a specific dysfunction of mitophagy in TECPR2 KO cells.

To test whether the primary dysfunction in mitophagy occurs due to inhibited mitochondrial delivery to the lysosomes, we tested several steps upstream of lysosomal consumption. As indicated in Fig. S3A, western blot analysis indicated stabilization of mitophagy factors PINK1 and Parkin in TECPR2 KO MEF cells. Consistently, confocal analysis revealed co-localization of PINK1 with mitochondria, marked by mitochondrial resident protein Cox4 (Fig. 3A). In addition, mitophagy receptor OPTN was also co-localized with mitochondrial Cox4 in KO cells to a higher degree than WT, suggesting elevated initiation of mitophagy (Fig. 3B). Finally, we examined the accumulation of mitochondrial cargo inside autophagic vesicles in TECPR2 KO cells by the Proteinase K protection assay^7^. The outer mitochondrial protein MFN2 was highly protected in TECPR2 KO cells and the autophagy and mitophagy receptor p62. In contrast, WT cells presented, as expected, low levels of protected mitochondrial cargo (Fig. 3C). This result suggests accumulation of mitochondria-containing autophagosomes in TECPR2 KO mice as observed by TEM analysis (Fig. 3D) and supported by elevated co-enrichment of autophagosome markers LC3B and p62 with mitochondrial protein VDAC under sucrose gradient fractionation of cellular membranes (Fig. S3B).

**Figure 3.**
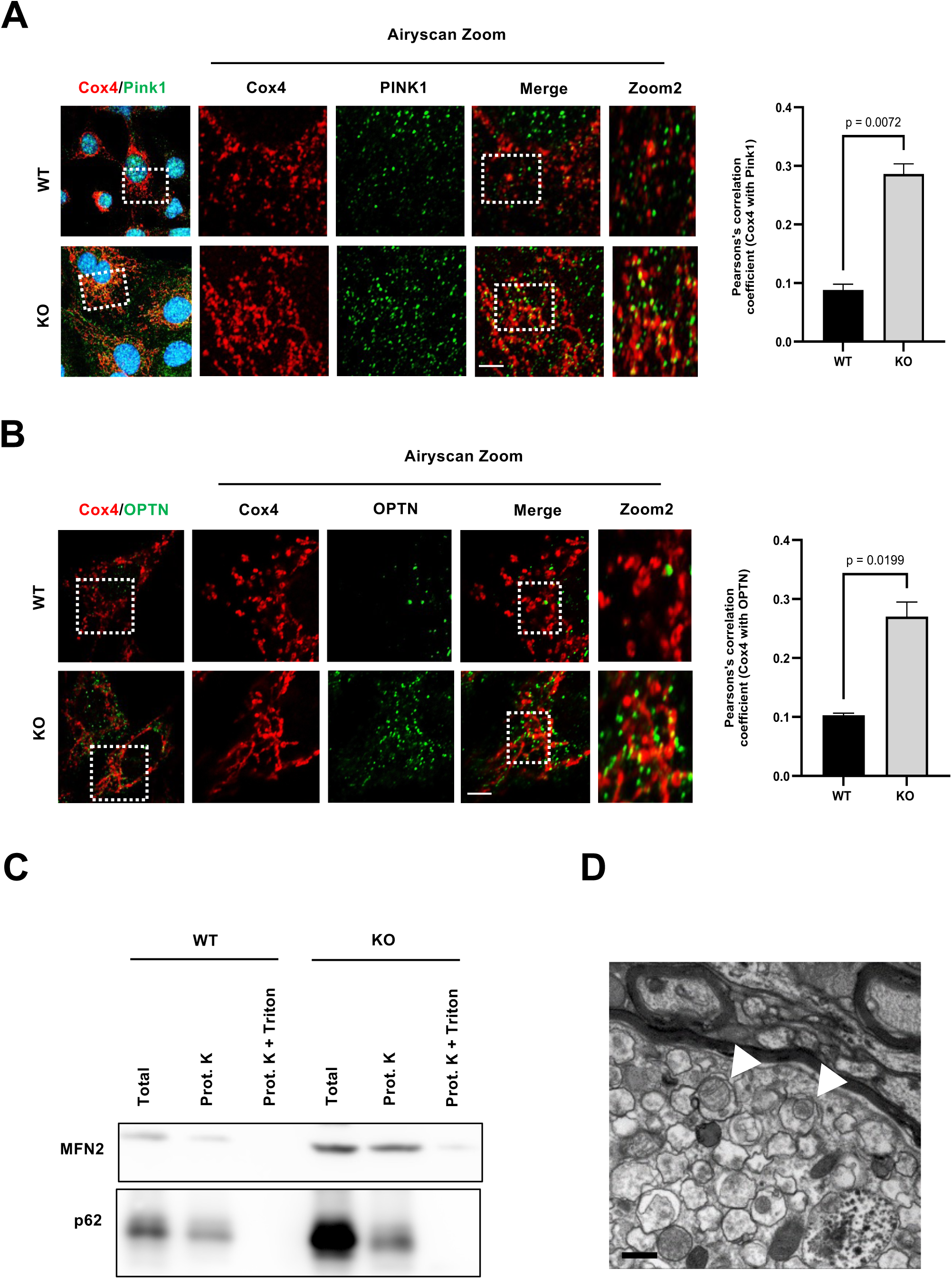
Elevated mitophagic activity in the absence of TECPR2. **A.** Immunofluorescence analysis of Cox4 and Pink1 stained in WT and TECPR2 KO cells, visualized by confocal microscopy in Airyscan settings for zoom-in images. Pearson’s colocalization analysis was performed, calculated, and presented (right panel) by three independent experiments: **p <= 0.01, determined by Student’s *t*-test. **B.** Immunofluorescence analysis of Cox4 and Optineurin stained in WT and TECPR2 KO cells, confocal microscopy in Airyscan settings for zoom-in images. Pearson’s colocalization analysis was performed, calculated, and presented (right panel) by three independent experiments: **p <= 0.01, determined by Student’s t-test. **C.** Autophagosome fractions from TECPR2 KO and WT MEF cells were subjected to Proteinase K and Triton X-100 treatments as described in Materials and Methods. **D.** TEM image indicates accumulation of double-membrane autophagic structures with mitochondrial cargo (white arrow) in a TECPR2 KO brain. Scale bar: 500 nm.

### ISR is activated in the absence of TECPR2

We next tested whether the mitochondrial dysfunction in TECPR2 KO cells leads to integrated stress response (ISR) through phosphorylation of eIF2α. As indicated in Fig. 4A, an increase in eIF2α phosphorylation was detected in TECPR2 KO MEF cells by immunoblotting with p-eIF2α and total eIF2α antibodies. Consistent with ISR induction^22^, increased protein and mRNA levels of Activating Transcription Factor 4 (ATF4) and Activating Transcription Factor 5 (ATF5) in TECPR2 KO MEF cells were detected (Fig. 4B-D). Moreover, immunofluorescence analysis revealed preferential nuclear localization of these transcription factors in TECPR2 KO cells (Fig. S4A and Fig. S4B), indicating activation of the mitochondrial unfolded protein response (UPR^mt^).

**Figure 4.**
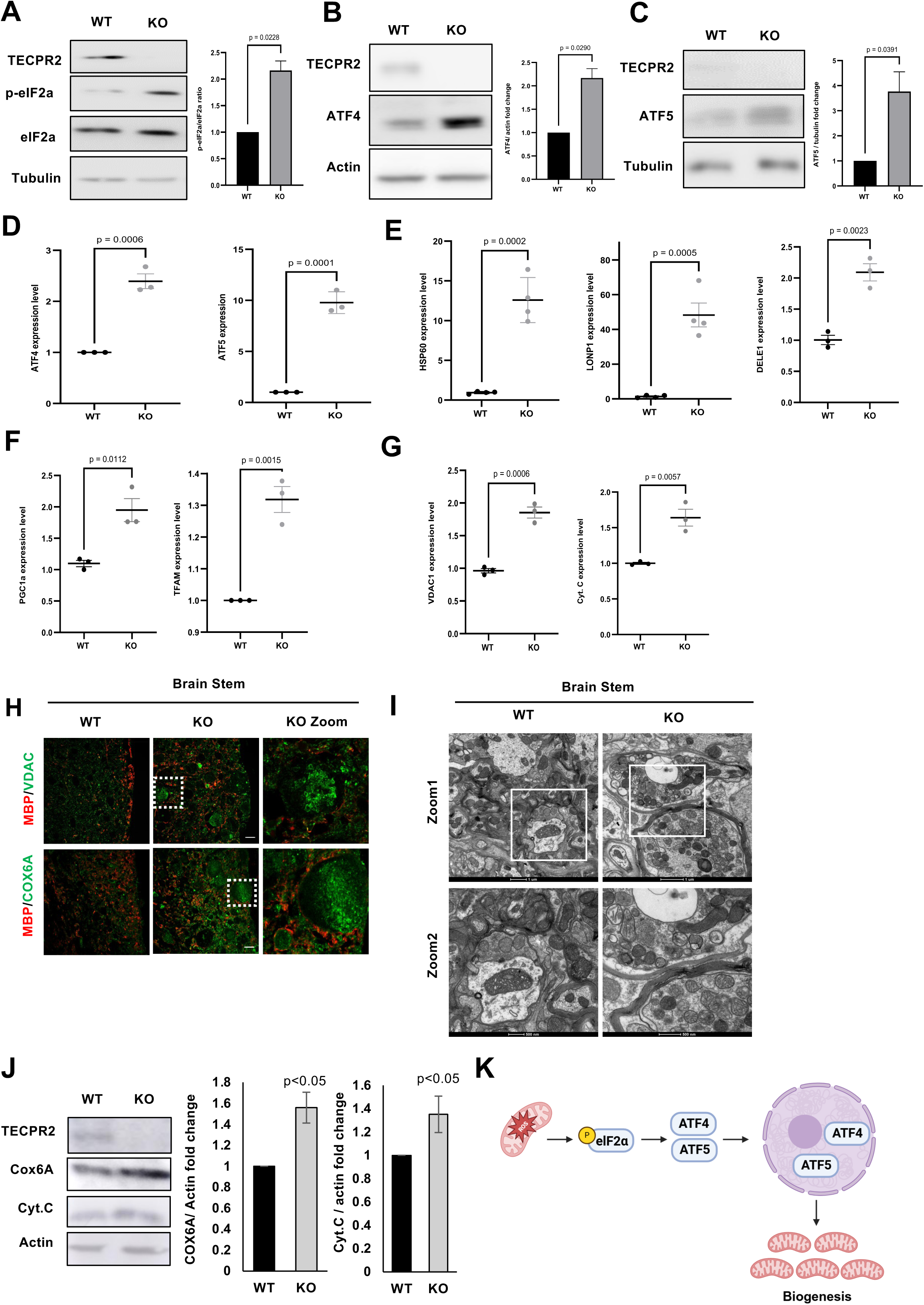
TECPR2 suppresses mitochondrial stress response. **A.** Cellular levels of peIF2α in WT and TECPR2 KO MEF cells were estimated by western blot analysis. The peIF2α/total eIF2α ratio was calculated in three independent experiments and presented as SEM, with student’s *t*-test *p < 0.05. **B.** Cellular levels of ATF4 in WT and TECPR2 KO MEF cells estimated by western blot analysis. **C.** Cellular levels of ATF5 in WT and TECPR2 KO MEF cells estimated by western blot analysis. **D.** Quantitative real-time PCR analysis for ATF4 and ATF5 expression in WT and TECPR2 KO MEF cells. **E.** Quantitative real-time PCR analysis for HSP60, LONP1, and DELE1 expressions in WT and TECPR2 KO MEF cells. **F.** Quantitative real-time PCR analysis for PGC1a and TFAM expression in WT and TECPR2 KO MEF cells. **G.** Quantitative real-time PCR analysis for VDAC1 and Cytochrome C expression in wildtype and TECPR2 KO MEF cells. Fold changes were calculated and presented with SEM of three independent experiments using Student’s *t*-test, *p < 0.05, **p < 0.01, and ***p < 0.001. **H.** Immunofluorescence analysis in spheroid regions of 9-month-old WT and TECPR2 KO mice stained with MBP for neuronal and VDAC or Cox6A for mitochondrial staining. **I.** TEM analysis in the brain stem region of 9-month-old WT and TECPR2 KO mice with accumulated mitochondrial content. **J.** Accumulation of mitochondrial content estimated by western blot analysis using mitochondrial markers Cox6A and Cytochrome C in WT and TECPR2 KO MEF cells. Protein levels of mitochondrial markers were calculated and presented with the SEM (n=3), *p < 0.05, determined by the student’s *t*-test. **K.** Schematic illustration depicting UPR^mt^ activation and elevated mitochondrial biogenesis in TECPR2 KO cells.

In line with activation of UPR^mt^ in TECPR2 KO cells, transcriptional levels of mitochondrial chaperonin Heat Shock Protein 60 (HSP 60), and mitochondrial Lon Protease 1 (LONP1), both playing a crucial role in maintaining mitochondrial integrity via protein folding and degradation^23,24^, HRI upstream activator, DAP3 Binding Cell Death Enhancer 1 (DELE1)^25^, were largely elevated (Fig. 4E). The knockdown of HRI led to partial reduction of p-eIF2α and ATF4 (Fig. S4G and Fig. S4H)

In light of the observed activation of UPR^mt^, we suspected an increased downstream mitochondrial biogenesis and accumulation of mitochondrial content in TECPR2 KO cells. To assess mitochondrial biogenesis, we tested the expression levels of peroxisome proliferator-activated receptor gamma coactivator 1-alpha (PGC1α), the master regulator of mitochondrial biogenesis^26^, and the mitochondrial transcription factor A (TFAM1), which maintains mitochondrial DNA copy number^27^. Indeed, expression of both genes was upregulated in TECPR2 KO cells (Fig. 4F). Additionally, we also tested the transcription and protein levels of several mitochondrial residents, such as mitochondrial outer membrane VDAC1 and matrix proteins, as cytochrome C (Cyt. C), Cox6A, Cox6B, and Cox6C, in MEFs and in human primary fibroblasts. Quantitative PCR (Fig. 4G and Fig. S4E) and western blot analyses (Fig. 4J and Fig. S4D) indicated elevation of these mitochondrial markers in TECPR2 KO cells. Moreover, the brain stem analyses of 9-month-old TECPR2 KO mice indicated significant accumulation of mitochondrial content visualized by confocal microscopy employing VDAC1 and COX6A antibodies (Fig. 4H). Mitochondrial accumulation in the brain stem of TECPR2 KO mice was also supported by TEM (Fig. 4I) and western blot analysis of the brain stem (Fig. S4F). Next, we visualized the mitochondrial network in WT and TECPR2 KO MEF cells using MitoTracker Green, which stains total mitochondrial content, and revealed an elevated number of elongated mitochondrial tubes accompanied by excessive network connectivity in TECPR2 KO cells (Fig. S4C).

Our results thus far indicate that the absence of TECPR2 leads to severe mitochondrial dysfunction accompanied by activation of ISR and UPR^mt,^ which is correlated with accumulation of mitochondrial content (Fig. 4K).

### The TECPR domain maintains mitochondrial homeostasis

To examine whether exogenous expression of TECPR2 may recover mitochondrial homeostasis in TECPR2 KO cells, we employed several stably expressing constructs of TECPR2 tagged with Halo-Flag tags as indicated in Fig. 5A. As depicted in Fig. 5B and Fig. 5C, expression of TECPR2 C-terminal domain led to suppression of mitochondrial ROS accumulation and partial recovery of mitochondrial membrane potential in a LIR-dependent manner. The transient expression of TECPR2 domains supported the relevance of the C-terminus for mitochondrial homeostasis, as the WD and middle domains of TECPR2 remained inert, while full-length TECPR2 and its C-terminus elevated mitochondrial membrane potential to a similar degree. Notably, the WT cells exhibited normal mitochondrial polarization after transient transfection of the indicated domains (Fig. S5A). The transcription levels of ATF4, ATF5, and LONP1 were consistently reduced upon C-terminal TECPR2 expression (Fig. S5B).

**Figure 5.**
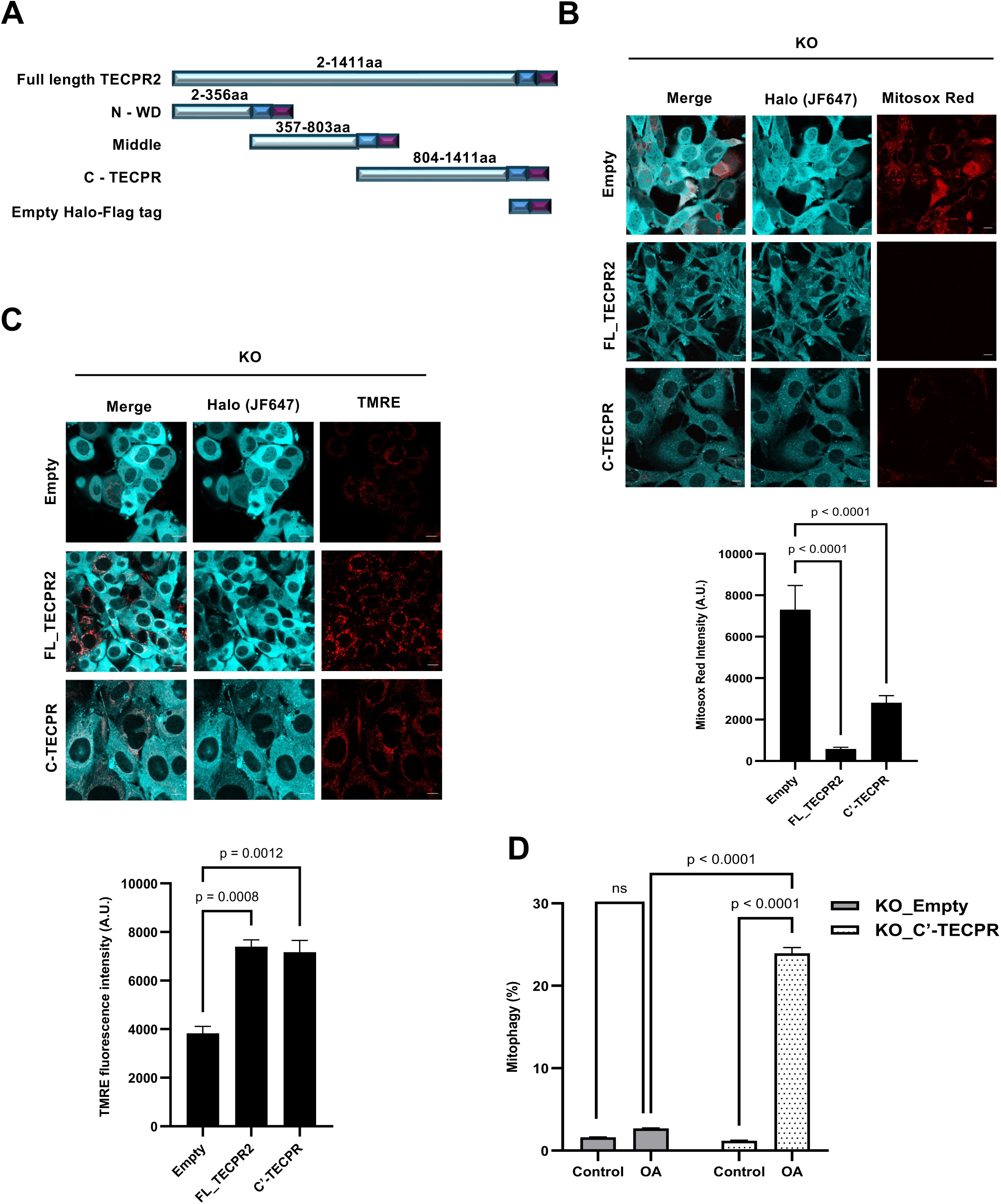
The TECPR domain maintains mitochondrial homeostasis. **A.** Schematic presentation of TECPR2 domains fused to Halo-tag and Flag-tag. **B.** Confocal microscopy visualization of mtROS stained with MitoSox Red in live MEF cells (TECPR2 KO background) stably expressing the following domains: Full-length TECPR2, C-terminal TECPR, and Empty-Halo-Flag constructs. The mean fluorescence intensity of MitoSox Red was calculated and presented with the SEM of three independent experiments, ****p < 0.0001, determined by one-way ANOVA with post-hoc Dunnett’s multiple comparison test. **C.** Confocal microscopy visualization of mitochondrial polarization by TMRE in live MEF cells (TECPR2 KO background) stably expressing the following domains: Full length TECPR2, C-terminal TECPR, and Empty-Halo-Flag constructs. The mean fluorescence intensity of TMRE was calculated and presented with the SEM of three independent experiments, **p < 0.01, ***p < 0.001, determined by one-way ANOVA with post-hoc Dunnett’s multiple comparison test. **D.** Flow cytometry of mitophagy flux analysis in MEF stably expressing Empty-Halo-Flag-tag cassette and C-terminal TECPR. Cells were treated with OA overnight. Data from three independent experiments; ****p < 0.0001; ns-non-significant using two-way ANOVA analysis, was done using Tukey’s multiple comparisons test.

To examine the effect of TECPR2 on mitophagy, we established cells stably expressing C-terminal TECPR2 domains on the background of mt-mKeima. Mitophagy was induced overnight by oligomycin/antimycin (OA), and flux was analyzed by flow cytometry analysis of mt-mKeima. As indicated in Fig. 5D and Fig. S5C, the C-terminal domain of TECPR2 recovered OA-induced mitophagy.

### ISR inhibition rescues mitochondrial function in TECPR2 mutants

Integrated stress response inhibitor (ISRIB) is a small molecule that inhibits phosphorylation of eIF2α, thus inhibiting ISR^28^. To test the role of ISR in mitochondrial phenotypes observed thus far in absence of TECPR2, we first treated WT and KO TECPR2 MEF cells with ISRIB and tested its effects by western blot analysis, where both eIF2α phosphorylation ratio and downstream UPR^mt^ sensors ATF4 and ATF5 levels were reduced for KO (Fig. 6A). Microscopic analysis further revealed translocation back to the cytosol of ATF4 and ATF5 in KO MEFs upon ISRIB treatment (Fig. S6A and Fig. S6B) and their transcriptional levels were also reduced (Fig. S6C). ISRIB treatment successfully recovered mitochondrial membrane potential in cell bodies and axons of KO DRG cells (Fig. 6B), and a similar effect was observed by both confocal microscopy and flow cytometry analysis in KO MEFs (Fig. S6E and Fig. S6F). Consistently, mtROS levels also were reduced in TECPR2 depleted cells upon ISRIB treatment (Fig. S6D). The transcriptional levels of mitochondrial biogenesis markers, PGC1α and TFAM1, mitochondrial quality control markers, LONP1 and HSP60, and general mitochondrial membrane markers VDAC1 and Cyt. C also significantly dropped upon treatment of KO cells with ISRIB (Fig. S6G).

**Figure 6.**
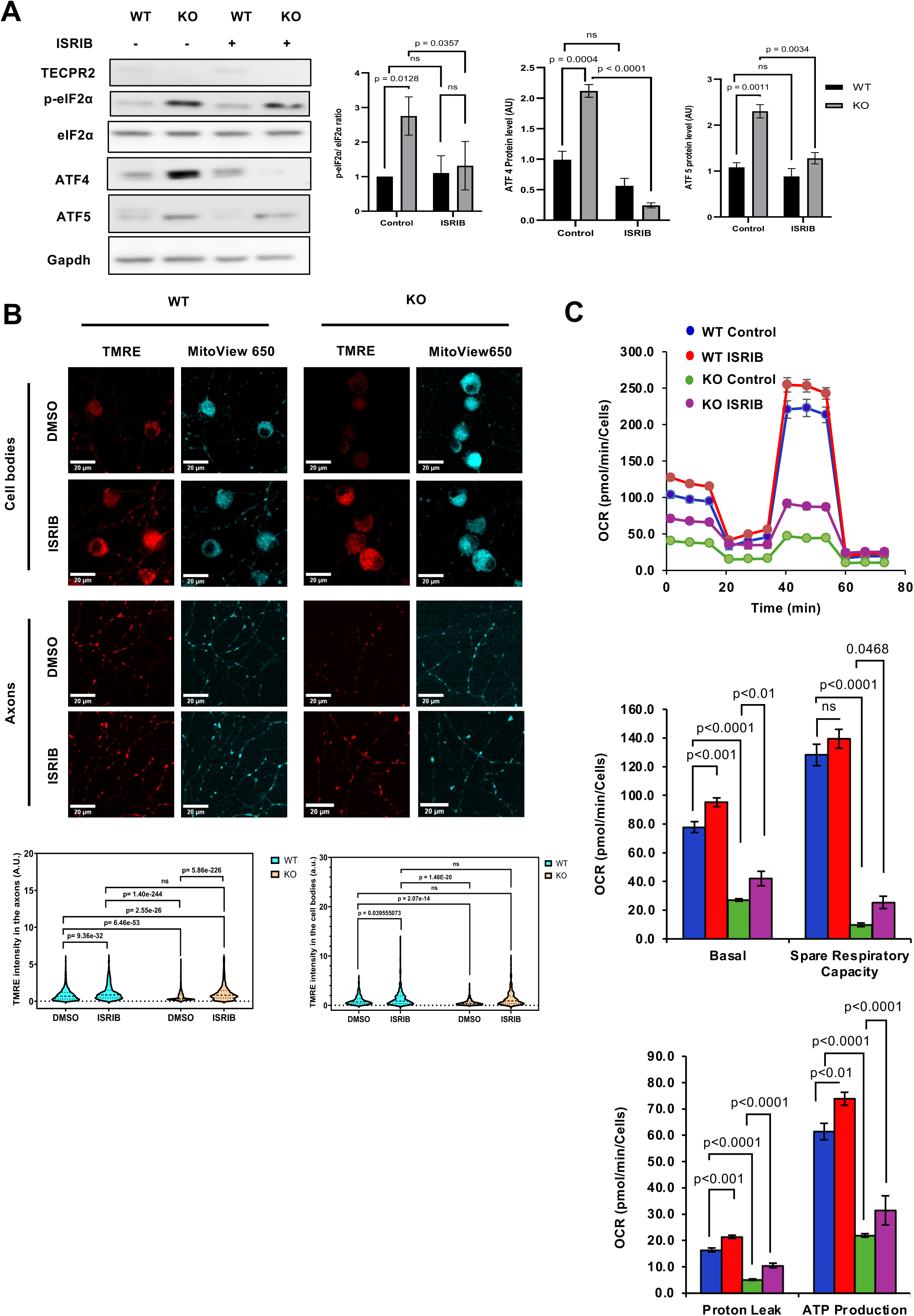
ISR inhibition rescues mitochondrial function in TECPR2 mutants. **A.** Western blot analysis of cell lysates isolated from TECPR2 KO and WT MEF cells treated with 200 nM ISRIB for 4 hours and immunoblotted for TECPR2, eIF2α (phospho and total), ATF5, and ATF4 using tubulin as a loading control. Data shown from three independent experiments. Significance was determined by two-way ANOVA followed by Tukey’s multiple comparison test. ****p < 0.0001, ***p < 0.001, **p < 0.001, *p < 0.05 and ns for non-significant. **B.** Confocal microscopy visualization of mitochondrial membrane potential by TMRE staining in cell bodies and axons of DRGs derived from WT and TECPR2 KO mice and treated with 200 nM ISRIB for 4 hours. Scale bar: 20 μm. Statistical analysis was conducted using the Kruskal–Wallis test followed by Dunn’s post-hoc test; *p < 0.05*, \*\*\*\**p < 0.0001, ns for non-significant. **C.** Seahorse analysis for quantification of mitochondrial respiration in wildtype and TECPR2 KO MEF cells treated with 200 nM ISRIB for 4 hours.

To further validate the rescue of mitochondrial well-being of TECPR2 KO cells by treatment with ISRIB, we estimated MEFs mitochondrial respiration by Seahorse mitostress assay, and observed recovery of oxidative phosphorylation and ATP production, hinting at a therapeutic potential for ISRIB in HSAN9 patients (Fig. 6C).

Finally, we examined whether ISRIB would also rescue OA-induced mitophagy flux in TECPR2 KO cells. Stably expressing mt-mKeima cells were pre-treated for 4h with ISRIB, followed by overnight treatment with OA, flow cytometry analysis revealed a significant induction in mitophagy flux in WT cells upon OA and OA+ISRIB treatments. However, OA-induced mitophagic flux in TECPR2 KO cells was not recovered by the ISRIB treatment (Fig. S6H).

## Discussion

TECPR2 has been implicated in multiple intracellular processes, including autophagy^1,6,7,16^, secretion^5^, and the endolysosomal system^29,30^. Here, we report a new link between aberrant mitophagy and activation of ISR in TECPR2 KO cells. This implicates TECPR2 in the regulation of mitochondrial homeostasis and elimination of defective mitochondria.

The dysfunction of mitochondria and impaired mitophagy are highly associated with neurodegenerative diseases, including Parkinson’s, Alzheimer’s, Huntington’s, amyotrophic lateral sclerosis (ALS), and others^13,15,31^. Hereditary spastic paraplegias (HSPs), are genetically heterogeneous neurodegenerative disorders linked to cellular and metabolic processes, including mitochondrial dysfunction and autophagy. HSPs were mainly associated with defects in endosome membrane trafficking, oxidative stress, and mitochondrial DNA polymorphisms^32–34^. Here we present evidence for a significant defect in mitochondrial well-being in cells derived from HSAN9 patients and from the HSAN9 mouse model^16^. The mitochondria in these cells exhibit reduced oxidative phosphorylation accompanied by accumulated mitochondrial ROS and membrane depolarization. All these phenotypes were recovered by expression of full-length TECPR2 or its C-terminal TECPR domain, supporting a role for TECPR2 in the maintenance of mitochondrial homeostasis. Moreover, we provide evidence for a beneficial effect of a pharmacological treatment with the ISR inhibitor ISRIB^35^, as well as other pharmaceutical and genetic ISR-inhibiting interventions, which lead to a partial recovery of the mitochondrial functions but not of mitophagy. Such an effect supports the notion that the mitochondrial defects observed in TECPR2-deficient cells are downstream of the inhibition of mitophagy.

ISRIB has shown efficacy in models of Alzheimer’s, Parkinson’s, and ALS, as well as in reversing age-related cognitive decline and improving memory in healthy animals^35^. Our findings provide the first indication of the potential of a beneficial effect of this drug in the HSAN9 model. The beneficial contribution here of inhibition of ISR, by itself a survival mechanism, is rather surprising; however, the reduction of mtROS observed in the presence of ISRIB may indicate a potential therapeutic strategy for HSAN9 and possibly for other HSP patients.

We have previously reported that TECPR2 is mostly required for autophagy taking place under basal growth conditions, hypothesizing that it may be required for selective autophagy^7^. The data presented in the current study indicate that TECPR2 is mostly required for mitophagy but not for other selective autophagy processes such as ER-phagy or pexophagy. Moreover, our finding that TECPR2 is needed for delivering autophagosomes carrying defective mitochondria to lysosomes is consistent with the recent report that TECPR1, its close homolog, is needed for the delivery of autophagosomes carrying aggregates to lysosomes^36^. Future studies are needed to determine whether TECPR2 participates in other selective autophagy processes and, more challenging, how it mediates targeting of specific sets of autophagosomes to lysosomes.

## Materials and Methods

### Study approval

Primary fibroblast cells were obtained from Sheba Medical Center in accordance with the Helsinki Declaration of 1975. All mouse work was performed according to the approved guidelines of the Institutional Animal Care and Use Committee at the Weizmann Institute of Science (WIS), IACUC approved numbers 10820514-2 and 28590716-3.

### Cell cultures and treatments

Primary fibroblast cells were grown on Dulbecco’s modified Eagle’s medium (DMEM; Gibco, 41965-039) supplemented with 20% fetal calf serum (FCS) and 1% L-Glutamine (Sigma, G5763) at 37°C in 5% CO_2_. MEF cells were grown on DMEM supplemented with 10% fetal calf serum (FBS; Invitrogen, 10270106) supplemented with 1% Sodium pyruvate (Gibco, 11,360,070) and 1% L-Glutamin at 37°C in 5% CO_2_. For induction of mitophagy, cells were treated with 20 µM Carbonyl cyanide 3-chlorophenylhydrazone CCCP (Sigma, C2759) for the indicated period. For ISR inhibition, ISRIB cells were treated with 200 nM ISRIB (Sigma, C2759) for 4 h or for the indicated period. PERK inhibitor II (GSK2656157) (Sigma-Aldrich, 504651), 1 µM for 4 h. For OA treatment, 10 µM oligomycin (Millipore Sigma, 495455) and 2.5 µM Antimycin a (Sigma-Aldrich, A8674) were combined for the indicated time. For mitochondrial ROS detection, cells were treated with 5 µM MitoSox Red, Mitochondrial superoxide indicator (Rhenium, M36008) for 15 min and analyzed by confocal analysis or by FACS. For antioxidant treatment, cells were incubated for 4 h with 10 mM N-Acetyl-L-cysteine NAC (Sigma Aldrich, A7250) or 5 µM Mitotempo (Sigma Aldrich, SML0737). Peroxisome proliferator 4-Phenylbutyric acid (Sigma-Aldrich P21005) 0.5 mM for 4 h and Deferiprone (Iron chelator) 1mM for 4 h. Lysosomal degradation was inhibited by 100 nM Bafilomycin A1 (LC Laboratories, B-1080) for 4 hours. All cell lines were routinely inspected for mycoplasma contamination monthly.

### Plasmids and siRNA

The cloning of full-length TECPR2 and its fragments into a Flag-labelled plasmid was performed using the restriction-free transfer PCR technique. The origin of this plasmid is pEGFP, where EGFP was replaced by a Flag tag. For *TECPR2* knockdown, cells were transfected with siRNA oligo *GUGCUGAGUUGGAAUGAAU* using DharmaPHECT reagent (Dharmacon, T-2001-03). For GCN2 knockdown, cells were transfected with ON-TARGETplus Mouse Eif2ak4 (27103) siRNA - SMARTpool (Dharmacon, L-044353-00-0005). The mt-mKeima plasmid (pHAGE-mt-mKeima) was purchased from Addgene (#131626). The CB5-mKeima plasmid for detecting ER-phagy was cloned in the Department of Life Sciences Core Facilities at the Weizmann Institute of Science. The SKL-mKeima, directed towards peroxisomes (pexo-mKeima) plasmid, was cloned in the Elazar lab using restriction-free cloning. GFP-Rab5Q79L (#28046) and GFP-Rab7Q67L (#169038), and mCherry-Parkin (#23956) plasmids were purchased from Addgene. MISSION esiRNA targeting mouse eIF2ak1 (siRNA HRI) (EMU061831) was purchased from Sigma Aldrich.

### Western blot analyses

Total cellular protein extracts were prepared in RIPA buffer [0.1 M NaCl; Bio-Lab Ltd, 21955 5 mM EDTA (J.T. Baker, 8993), 0.1 M sodium phosphate, pH 7.5; Sigma, 342483, 1% Triton X-100 (Sigma, X100), 0.5% sodium deoxycholate (Sigma, D6750), 0.1% sodium dodecyl sulfate (Sigma, L4509)] with a protease inhibitor cocktail (PIC; Merck, 539134). The extracts were centrifuged at 16,000 x g for 15 min at 4°C, and protein concentrations were determined using Bio-Rad Protein Assay Dye Reagent Concentrate (Bio-Rad, 500-0006). Total proteins (30 µg) were separated by SDS−PAGE (12% polyacrylamide) and transferred to a nitrocellulose membrane (Bio-Rad, 1704159). The membrane was blocked in phosphate-buffered saline (PBS) with 5% skim milk for 1 h at room temperature, and then incubated with the appropriate primary antibody overnight at 4°C. It was then washed three times with PBS-TWEEN 20 (0.1%, Sigma, P1379) and incubated with the secondary antibody (goat anti-mouse or goat anti-rabbit) for 1 h at room temperature. Finally, the membrane was washed three times and specific proteins were visualized using the Enhanced Chemiluminescence (ECL) detection system (Biological Industries, 20-500-120). The antibodies employed for this assay are described in Table 1.

**Table 1.** Antibodies used in this study.

| **Antibody** | **Host** | **Cat. Number** | **Source** | **Working Concentration** |
| --- | --- | --- | --- | --- |
| TECPR2 | Rabbit | Custom | Provided by Prof. Behrends, LMU, München | 1:1000 WB |
| LC3B | Rabbit | Custom | Custom-approval made (our lab) | 1:1000 WB<br>1:200 IF |
| SQSTM1/p62 | Mouse | H00008878-M01 | Abnova | 1:3000 WB<br>1:200 IF |
| Actin | Mouse | MAB1501 | Sigma | 1:5000 WB |
| Flag | Mouse | F1804 | Sigma | 1:2000 WB |
| MBP | Rat | Ab7349 | Abcam, UK | 1:250 IF |
| Cox4 | Mouse | MAB6980 | R&D Systems | 1:200 IF |
| Cox6A1 | Rabbit | Ab236889 | Zotal | 1:1000 WB<br>1:200 IF |
| VDAC | Rabbit | CST-4661S | Cell Signaling Technology | 1:1000 WB<br>1:200 IF |
| Cytochrome C | Mouse | Ab110325 | Zotal | 1:1000 WB |
| PINK1 | Rabbit | BC100-494 | Biotest Ltd. | 1:1000 WB<br>1:200 IF |
| Parkin | Mouse | Ab77924 | Zotal | 1:1000 WB |
| OPTN | Rabbit | Ab213556 | Zotal | 1:1000 WB<br>1:200 IF |
| Tubulin | Mouse | T9026 | Sigma | 1:3000 WB |
| ATF4 | Rabbit | CST-11815S | Cell Signaling Technology | 1:1000 WB<br>1:250 IF |
| ATF5 | Rabbit | EPR18286 | Abcam | 1:1000 WB<br>1:250 IF |
| Phospho-eIF2α (Ser51) (D9G8) | Rabbit | CST-3398 | Cell Signaling Technology | 1:1000 WB |
| Total eIF2α | Goat | AF3997-SP | Biotest Ltd. | 1:1000 WB |
| MFN2 antibody | Mouse | H00009927-M03 | Abnova | 1:1000 WB |
| GAPDH | Mouse | MAB374 | Merck | 1:5000 WB |

IF, immunofluorescence; DSHB, Developmental Studies Hybridoma Bank; WB, western blot

### Membrane flotation assay

Cells were homogenized with a Balch homogenizer. Homogenates (2mg protein) were adjusted to 2M sucrose, placed at the bottom of rotor tubes, overlaid with 1.75, 1.5, 1.25, 1, and 0.75M sucrose, and centrifuged in the SW-28 rotor (Beckman) at 28,000 rpm (slow acceleration and deceleration) overnight, 4°C. Fractions from the top of the gradient were collected, and sucrose densities were estimated from their refractive indices. Proteins from each fraction (180 µl) were precipitated by 10% trichloroacetic acid (TCA; Merck, T6399), washed with 100% cold acetone, boiled in sample buffer, and immunoblotted.

### Proteinase K protection assay

Organs or cells cultured and treated in 15 cm dishes were detached by trypsin, washed (PBS x 3), resuspended in 4 pellet volumes of homogenization buffer (10 mM Tris, pH 7.4, and 0.25M sucrose) supplemented by protease inhibitors, and homogenized on ice with a Balch homogenizer. Unbroken cells and nuclei were removed (700 x g, 5 min, 4°C), and equal amounts of homogenate were centrifuged using a TLA 120.2 rotor (90,000 rpm, 30 min, 4°C) for cytosol and membrane fractions. The pellets were resuspended in homogenization buffer. Each fraction was then divided into equal volumes for treatment with (30 min, 37°C) proteinase K (10 µg/ml, Merck, 1245680) or Triton X-100 (0.4% (v/v)) with proteinase K, as indicated. Treatments were terminated by the addition of phenylmethylsulfonyl fluoride (PMSF; Sigma, 78830) (200 mM, 10 min on ice), and proteins were precipitated by 10% TCA, boiled in sample buffer, and immunoblotted.

### Immunostaining

Cells cultured on sterile coverslips (13 mm) and treated as indicated were fixed and permeabilized by 100% methanol (Bio-Lab Ltd, 136805) for 5 min at−20°C, washed three times with PBS, blocked (10% FCS in PBS, 30 min at room temperature), and incubated with primary antibody (1 h at room temperature or Overnight at 4°C), washed (as above) and incubated with secondary antibody (45 min at room temperature). We used methanol fixation for Cox4-Pink1, Cox4-OPTN, Cox4-LC3B and mCherry parkin-Cox4 and LC3B staining. For other IFA experiments (ATF4, ATF5, Cox4, and ATF5), we fixed cells with 4% paraformaldehyde for 15 min at room temperature. Permeabilization with 0.1% permeabilization buffer (0.1% Triton X-100 in PBA) washed two times with PBS, blocked (10% FCS in PBS, 30 min at room temperature), and incubated with primary antibody (1 h at room temperature or overnight at 4°C), washed (as above) and incubated with secondary antibody (45 min at room temperature).

### Imaging

Wide-field images were obtained with an Axio Observer 7 inverted microscope system using an objective Plan-Apochromat 20×/0.8 M27 and 1.6× tube lens. Images were analyzed using ZEN Blue software (Zeiss) and ImageJ. Confocal images were obtained with a Zeiss LSM 880 Axio Observer 7 laser scanning confocal microscope using Plan-Apochromat 20×/0.8 M27 or alpha Plan-Apochromat 100×/1.46 Oil DIC M27 Elyra objectives. Images were analyzed using Zen Blue software (Zeiss) and ImageJ. Histological images were obtained with a pco.edge 4.2 4MP camera using a Plan-Apochromat 20× objective and PanoramA SCAN 150 software. Images were analyzed using CaseViewer 2.2 software (3DHISTECH) and ImageJ.

### Transmission electron microscopy

Samples were fixed with 4% PFA, 2% glutaraldehyde, in 0.1 M cacodylate buffer containing 5 mM CaCl_2_ (pH 7.4), post-fixed in 1% osmium tetroxide supplemented with 0.5% potassium hexacyanoferrate trihydrate and potassium dichromate in 0.1 M cacodylate for 1 h, stained with 2% uranyl acetate in double-distilled water for 1 h, dehydrated in graded ethanol solutions, and embedded in epoxy resin (Agar Scientific, UK). Ultrathin sections (70-90 nm), obtained with a Leica Ultracut UCT microtome, were stained with lead citrate and then examined using either a Phillips CM-12 transmission electron microscope (TEM) equipped with a Gatan One View camera, or an FEI Tecnai SPIRIT TEM equipped with a bottom-mounted 2k × 2k FEI Eagle CCD camera.

### RNA extraction and real-time PCR

Total cell RNA was extracted with NucleoSpin RNA kit (Bioanalysis, 020009764). Reverse transcription was performed using qScript cDNA Synthesis Kit (Quantabio, 95047-100). Real-time quantitative PCR (qPCR) was performed with Fast SYBR™ Green Master Mix (Applied Biosystems, 4385612) according to the manufacturer’s instructions on the Viia - 7 and Quant Studio ABI (Applied Biosystems). Primers are used against cDNA encoding genes, as shown in Table 2.

**Table 2.**
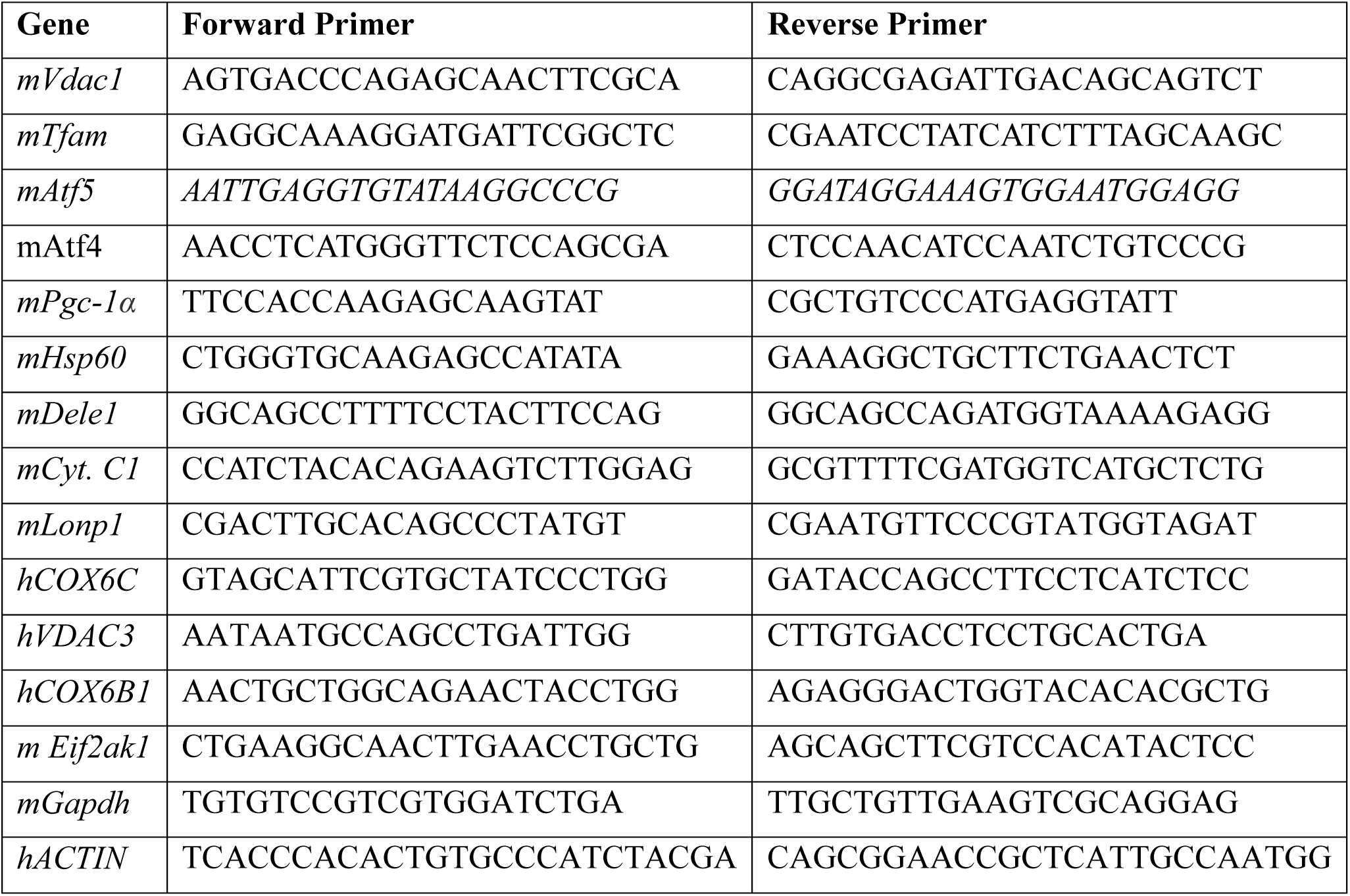
Real-time PCR primers used in this study.

### Seahorse

The key parameters of mitochondrial respiration were measured by Cell Mito Stress Test in primary human fibroblast cells derived from a healthy individual and a SPG49 patient or MEF cells derived from WT and *TECPR2* KO mice employing a Seahorse XFe96 as detailed in Gu et al.^37^.

### Flow cytometry analysis

According to manufacturer instructions, cells were labeled with MitoSOX Red or Tetramethylrhodamine Ethyl Ester Perchlorate TMRE (Invitrogen, T669) reagents. Labeled cells were analyzed on ZE5 cell analyzer (Bio-Rad Laboratories). Excitation of MitoSOX Red was done with a 561nm laser line, while emitted light was collected through a 593/52nm bandpass. Excitation of TMRE was done with a 561nm laser line, and emitted light was collected through a 577/15nm bandpass filter.

### Mitophagy analysis (confocal microscopy and flow cytometry)

For Live confocal assessment of mitophagy, cells were treated with 20 µM CCCP or 10 µM Oligomycin + 2.5 µM Antimycin A for 4 h, followed by confocal imaging of live MEF cells stably expressing mt-mKeima was performed using an inverted Leica SP8 STED3× microscope, equipped with internal Hybrid detectors and acousto-optical tunable filter (Leica Microsystems CMS GmbH, Germany), a white light laser excitation laser (WLL), and an Argon laser. Imaging was done with an HC PL APO 86×/1.20 water STED white objective, a numerical aperture of 1.2. Cells expressing mt-mKeima were excited in two sequences, first with 10% 458 nm with a line of an Argon laser, and second at 10% 586 nm with the WLL. mt-mKeima emission signal was collected at 596 to 680 nm in both sequences. Images were acquired using the galvometric scanner in the format 872×872 with a line average of 4 at a scan speed of 600 Hz and a pinhole of 1 airy unit with 16-bit depth. The acquired images were visualized using LAS X software (Leica Application Suite X, Leica Microsystems CMS GmbH). **Mitophagy analysis by flow cytometry:** Cells were treated overnight with 20 µM CCCP or 10 µM Oligomycin + 2.5 µM Antimycin A for overnight treatment in the presence of 10 µM Q-Vd-OPh-hydrate (Enco Scientific Services, A14915). PH-dependent fluorescent protein mt-mKeima was excited with a 561nm laser line (for acidic environment) or a 405nm laser line (for physiological pH). At the same time, emitted light was collected through 615/24 bandpass filters and analyzed using a ZE5 cell analyzer (Bio-Rad Laboratories). Analysis was performed on FCS Express 7 Software (De Novo Software, Glendale, CA).

### ERphagy analysis

For Live confocal assessment of ERphagy, cells were treated with 1 µM tunicamycin for 4 hours, followed by Confocal imaging of live MEF cells stably expressing Cb5-mKeima, which was performed using a Zeiss LSM 880 Axio Observer Confocal microscope. Cells expressing Cb5-mKeima were excited in two sequences, first with 10% 458 nm with an Argon laser and second at 5% 561 nm with the WLL. Cb5-mKeima emission signal was collected and a pinhole of 1 airy unit with 16-bit depth. The acquired images were visualized using Zen Blue software. **ERphagy analysis by flow cytometry:** Cells were treated with 1 µM tunicamycin overnight. Excitation of pH-dependent fluorescent protein Cb5-mKeima was done with a 561nm laser line (for acidic environment) or with a 405nm laser line (for physiological pH), while emitted light was collected through 615/24 bandpass filters and analyzed on a ZE5 cell analyzer. Analysis was performed on FCS Express 7 Software (De Novo Software, Glendale, CA).

### Pexophagy analysis

For live confocal assessment of pexophagy, cells were treated with 1mM DFP for 24 h, followed by confocal imaging of live MEF cells stably expressing SKL-mKeima, which was performed using a Zeiss LSM 880 Axio Observer Confocal microscope. Cells expressing SKL-mKeima were excited in two sequences, first with 10% 458 nm with an Argon laser and second at 5% 561 nm with the WLL. Pexo-mKeima emission signal was collected, and a pinhole of 1 airy unit with 16-bit depth. The acquired images were visualized using Zen Blue software. **Pexophagy analysis by flow cytometry:** Cells were treated with 1 mM DFP for 24 h. Excitation of pH-dependent fluorescent protein pexo-mKeima was done with a 561nm laser line (for acidic environment) or with a 405nm laser line (for physiological pH), while emitted light was collected through 615/24 bandpass filters and analyzed on a ZE5 cell analyzer. Analysis was performed on FCS Express 7 Software (De Novo Software, Glendale, CA).

### Statistical analysis

Where appropriate, statistical significance between data sets was analyzed by *t*-tests, Analysis of variance one-way (Anova), two-way (Anova), and Kruskal–Wallis test, followed by Dunnet’s post-hoc test using GraphPad Prism. n.s. for non-significant; *p < 0.05, **p < 0.01, ***p < 0.001, ****p < 0.0001.

## Acknowledgments

Z.E. is the Harold Korda Chair of Biology incumbent, supported by the Molly and Steven Elias Foundation and the Richard F. Goodman Yale/Weizmann Exchange Program. We are grateful for funding from the Israel Science Foundation (Grant #1272/24), the U.S.-Israel Binational Science Foundation (Grant #2023211), the Weizmann - Gladys Monroy and Larry Marks Center for Brain Disorders, and the Azrieli Institute for Brain and Neural Sciences Research Centers.

Part of the images in this paper were acquired at the Advanced Optical Imaging Unit, de Picciotto-Lesser Cell Observatory unit at the Moross Integrated Cancer Center Life Science Core Facilities, Weizmann Institute of Science.

## Author Contributions

1. M. C., M. F., N. S., O. S., K. K., O. G., B. C. T. Y., S. F., A. D., experimental and intellectual contribution, E. K., flow cytometry, I. G., confocal microscopy imaging, S. C., E. A. Seahorse analyses, Y. P. cloning, N. D. transmission microscopy, Z. E. principal investigator

## Ethics Declaration

The authors declare no competing interests.

## Data availability statement

The authors confirm that the data supporting the findings of this study are available within the article and its supplementary materials.

## Notes

### Competing Interest Statement

The authors have declared no competing interest.

### Summary of Updates

Just fixing the authors' list order - moving Zvulun Elazar to the last place to fit with the actual PDF.

## References

1 Oz-Levi, D. et al. Mutation in TECPR2 reveals a role for autophagy in hereditary spastic paraparesis. Am J Hum Genet 91, 1065–1072, (2012).

2 Oz-Levi, D., Gelman, A., Elazar, Z. & Lancet, D. TECPR2. Autophagy 9, 801–802, (2013).

3 Guan, Y. et al. Novel detection of mutation in the TECPR2 gene in a Chinese hereditary spastic paraplegia 49 patient: a case report. BMC Neurology 22, (2022).

4 Heimer, G. et al. TECPR2 mutations cause a new subtype of familial dysautonomia like hereditary sensory autonomic neuropathy with intellectual disability. Eur J Paediatr Neurol 20, 69–79, (2016).

5 Stadel, D. et al. TECPR2 Cooperates with LC3C to Regulate COPII-Dependent ER Export. Mol Cell 60, 89–104, (2015).

6 Behrends, C., Sowa, M. E., Gygi, S. P. & Harper, J. W. Network organization of the human autophagy system. Nature 466, 68–76, (2010).

7 Fraiberg, M. et al. Lysosomal targeting of autophagosomes by the TECPR domain of TECPR2. Autophagy 17, 3096–3108, (2021).

8 Palikaras, K., Lionaki, E. & Tavernarakis, N. Mechanisms of mitophagy in cellular homeostasis, physiology and pathology. Nature Cell Biology 20, 1013–1022, (2018).

9 Onishi, M., Yamano, K., Sato, M., Matsuda, N. & Okamoto, K. Molecular mechanisms and physiological functions of mitophagy. The EMBO Journal 40, (2021).

10 Baker, B. M., Nargund, A. M., Sun, T. & Haynes, C. M. Protective coupling of mitochondrial function and protein synthesis via the eIF2alpha kinase GCN-2. PLoS Genet 8, e1002760, (2012).

11 Chakrabarty, Y., Yang, Z., Chen, H. & Chan, D. C. The HRI branch of the integrated stress response selectively triggers mitophagy. Molecular Cell 84, 1090–1100.e1096, (2024).

12 Perea, V. et al. Pharmacologic activation of a compensatory integrated stress response kinase promotes mitochondrial remodeling in PERK-deficient cells. Cell Chemical Biology 30, 1571–1584.e1575, (2023).

13 Dikic, I. & Elazar, Z. Mechanism and medical implications of mammalian autophagy. Nat Rev Mol Cell Biol 19, 349–364, (2018).

14 Mizushima, N. & Levine, B. Autophagy in Human Diseases. N Engl J Med 383, 1564–1576, (2020).

15 Fraiberg, M. & Elazar, Z. Genetic defects of autophagy linked to disease. Prog Mol Biol Transl Sci 172, 293–323, (2020).

16 Tamim-Yecheskel, B. C. et al. A tecpr2 knockout mouse exhibits age-dependent neuroaxonal dystrophy associated with autophagosome accumulation. Autophagy 17, 3082–3095, (2021).

17 Leung, D. T. H. & Chu, S. Measurement of Oxidative Stress: Mitochondrial Function Using the Seahorse System. Methods Mol Biol 1710, 285–293, (2018).

18 Narendra, D. P. & Youle, R. J. The role of PINK1-Parkin in mitochondrial quality control. Nat Cell Biol 26, 1639–1651, (2024).

19 Katayama, H., Kogure, T., Mizushima, N., Yoshimori, T. & Miyawaki, A. A sensitive and quantitative technique for detecting autophagic events based on lysosomal delivery. Chem Biol 18, 1042–1052, (2011).

20 Sun, N. et al. A fluorescence-based imaging method to measure in vitro and in vivo mitophagy using mt-Keima. Nature Protocols 12, 1576–1587, (2017).

21 Wang, C. A Sensitive and Quantitative mKeima Assay for Mitophagy via FACS. Current Protocols in Cell Biology 86, (2020).

22 Melber, A. & Haynes, C. M. UPR(mt) regulation and output: a stress response mediated by mitochondrial-nuclear communication. Cell Res 28, 281–295, (2018).

23 Singh, M. K. et al. Molecular Chaperonin HSP60: Current Understanding and Future Prospects. Int J Mol Sci 25, (2024).

24 Ngo, J. K. & Davies, K. J. Importance of the lon protease in mitochondrial maintenance and the significance of declining lon in aging. Ann N Y Acad Sci 1119, 78–87, (2007).

25 Sekine, Y. et al. A mitochondrial iron-responsive pathway regulated by DELE1. Mol Cell 83, 2059–2076 e2056, (2023).

26 Schreiber, S. N. et al. The estrogen-related receptor alpha (ERRalpha) functions in PPARgamma coactivator 1alpha (PGC-1alpha)-induced mitochondrial biogenesis. Proc Natl Acad Sci U S A 101, 6472–6477, (2004).

27 Ekstrand, M. I. et al. Mitochondrial transcription factor A regulates mtDNA copy number in mammals. Hum Mol Genet 13, 935–944, (2004).

28 Coulson, R. L., et al. Translational modulator ISRIB alleviates synaptic and behavioral phenotypes in Fragile X syndrome. iScience 27, 109259, (2024).

29 Nalbach, K. et al. Spatial proteomics reveals secretory pathway disturbances caused by neuropathy-associated TECPR2. Nat Commun 14, 870, (2023).

30 Paul, S. et al. TECPR2 is a Rab5 effector that regulates endosomal cargo recycling (Cold Spring Harbor Laboratory, 2024).

31 Yan, X., Wang, B., Hu, Y., Wang, S. & Zhang, X. Abnormal Mitochondrial Quality Control in Neurodegenerative Diseases. Front Cell Neurosci 14, 138, (2020).

32 Awuah, W. A., et al. Hereditary spastic paraplegia: Novel insights into the pathogenesis and management. SAGE Open Med 12, 20503121231221941, (2024).

33 Pfeffer, G. et al. Mutations in the SPG7 gene cause chronic progressive external ophthalmoplegia through disordered mitochondrial DNA maintenance. Brain 137, 1323–1336, (2014).

34 Wedding, I. M. et al. Spastic paraplegia type 7 is associated with multiple mitochondrial DNA deletions. PLoS One 9, e86340, (2014).

35 Costa-Mattioli, M. & Walter, P. The integrated stress response: From mechanism to disease. Science 368, (2020).

36 Wetzel, L. et al. TECPR1 promotes aggrephagy by direct recruitment of LC3C autophagosomes to lysosomes. Nat Commun 11, 2993, (2020).

37 Gu, X., Ma, Y., Liu, Y. & Wan, Q. Measurement of mitochondrial respiration in adherent cells by Seahorse XF96 Cell Mito Stress Test. STAR Protoc 2, 100245, (2021).

