## Supplemental Figures 1 to 6 for "TECPR2 maintains mitochondrial homeostasis in neurodegeneration"

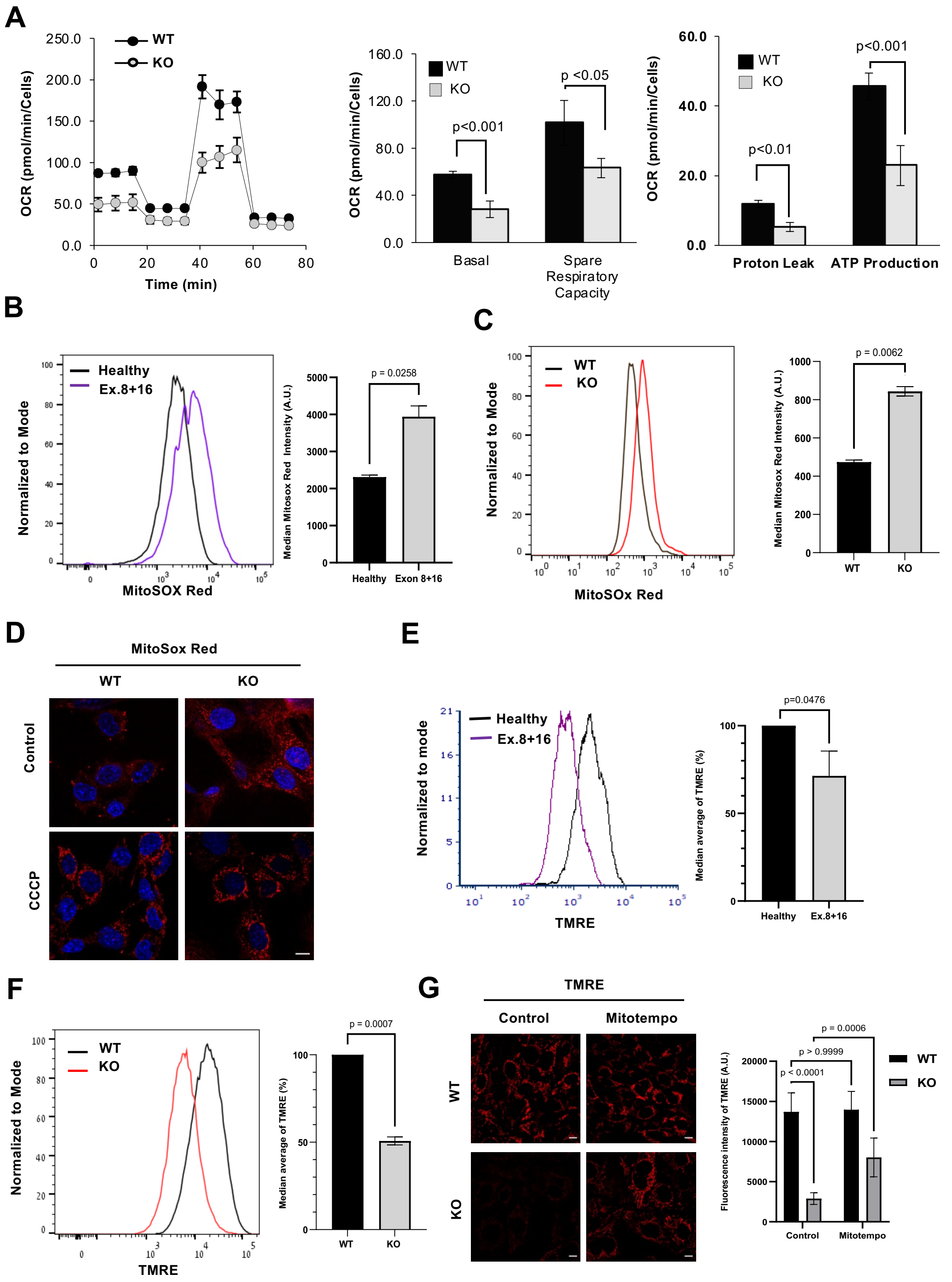


**Figure S1. TECPR2 maintains mitochondrial homeostasis. A.** Seahorse analysis for quantification of mitochondrial respiration in MEF cells obtained from the TECPR2 KO mouse model. **B.** Flow cytometry analysis of MitoSox Red in HSAN9 patient-derived fibroblasts versus healthy control, and **C.** MEF cells derived from WT and TECPR2 KO mice. The median intensity of Mitosox Red was calculated and presented with the SEM of three independent experiments, *p < 0.05; **p < 0.01 determined by Student's *t*-test. **D.** Confocal microscopy visualization of mtROS in CCCP-treated WT and TECPR2 KO MEF cells. Scale bar: 20 μm. **E.** Flow cytometry analysis for mitochondrial membrane potential in HSAN9 patient-derived fibroblasts versus healthy controls and in **F.** MEF cells derived from WT and TECPR2 KO mice. The median intensity of TMRE was calculated and presented with the SEM of three independent experiments, *p < 0.05; ***p < 0.001 determined by Student's *t*-test. **G.** Confocal microscopy visualization of mitochondrial membrane potential recovery indicated by TMRE staining in TECPR2 KO MEF cells treated with mtROS scavenger, MitoTempo.

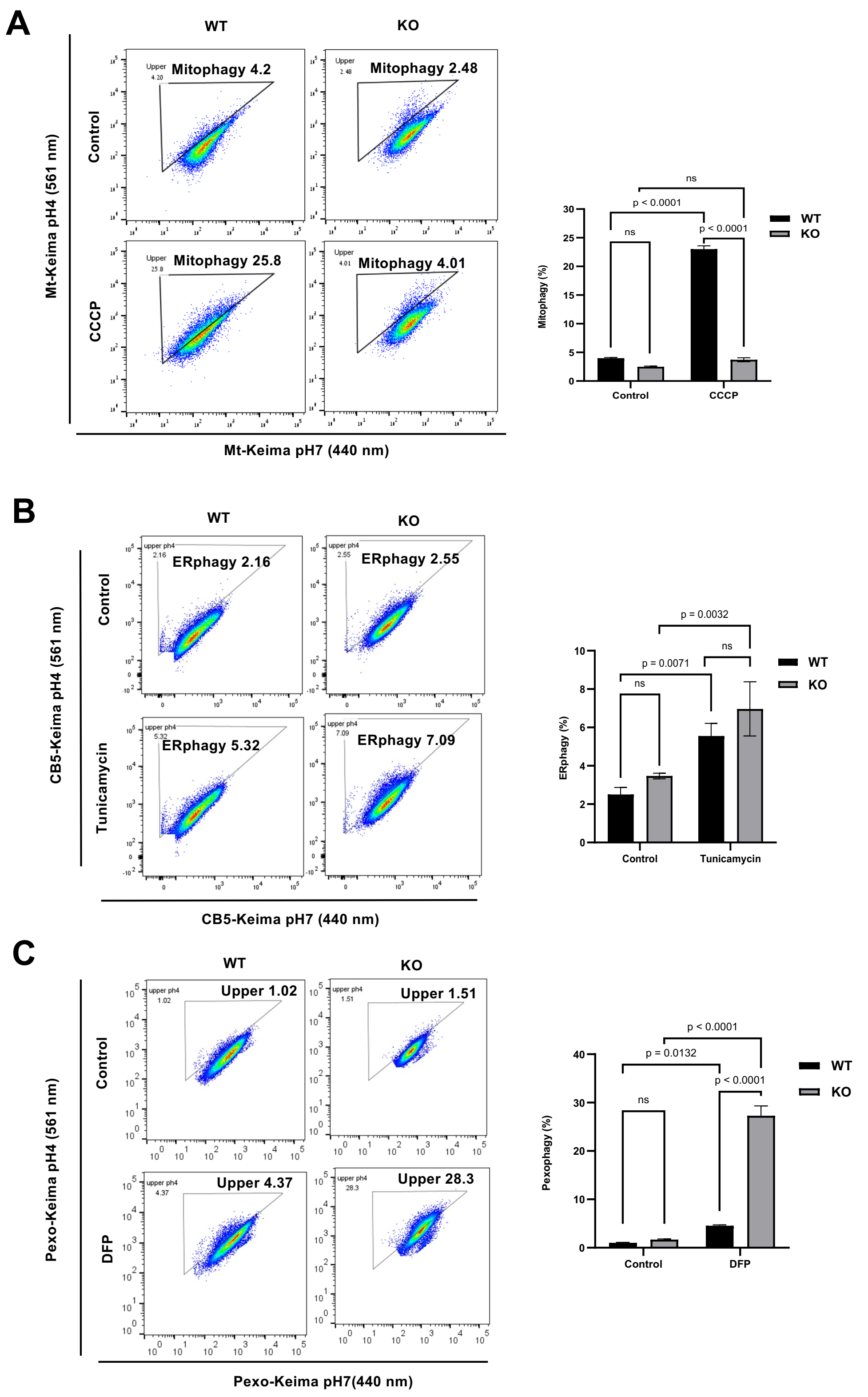


**Figure S2. TECPR2 supports mitophagic flux upon mitochondrial stress. A.** Flow cytometry analysis of mitophagy in mKeima stable WT and TECPR2 KO cells treated with

CCCP for 16 h. Data is shown from three independent biological replicates. ***$p < 0.001$, analyzed by using two-way ANOVA. **B.** Flow cytometry analysis of ERphagy in Cb5-Keima stable TECPR2 KO and WT cells treated with tunicamycin for 16 h. Data were summarized from three independent replicates. **$p < 0.01$ between WT control and WT tunicamycin, **$p < 0.01$ between KO Control and KO tunicamycin, analyzed by two-way ANOVA. **C.** FACS analysis of pexo-phagy in SKL-Keima (pexo-keima) stable TECPR2 KO, and WT cells treated with DFP for 24 h. Data was summarized from three independent replicates and analysed using two-way ANOVA with Tukey's multiple comparison test. ****$p < 0.0001$, *$p < 0.05$ and ns, nonsignificant.

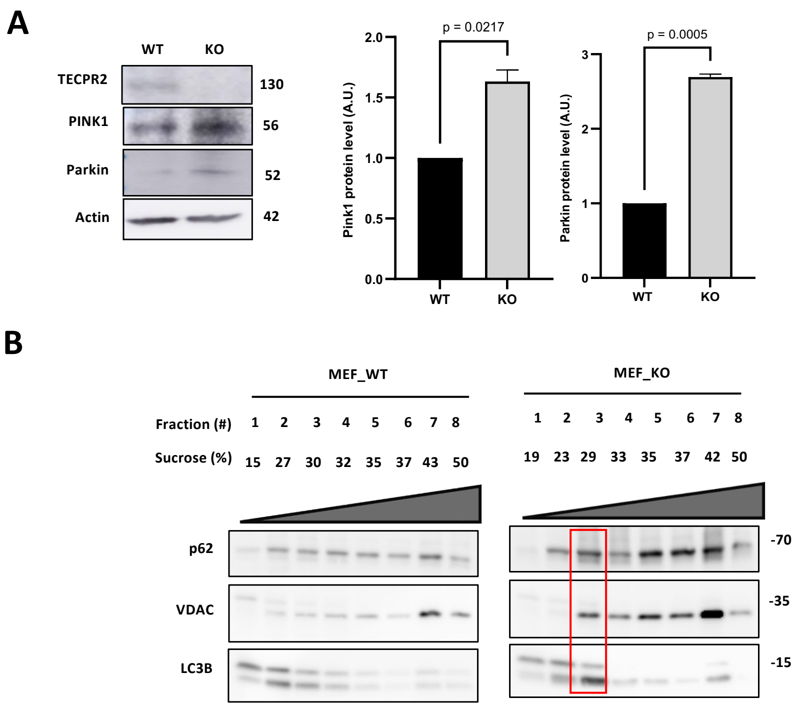


**Figure S3. Elevated mitophagic activity in the absence of TECPR2. A.** Accumulation of PINK and Parkin in TECPR2 KO cells indicated by western blot analysis. **B.** Homogenates of TECPR2 KO and WT MEF cells were floated over a sucrose gradient as described in Materials and Methods, and the proteins of the collected fraction were immunoblotted for p62, LC3B, and VDAC.

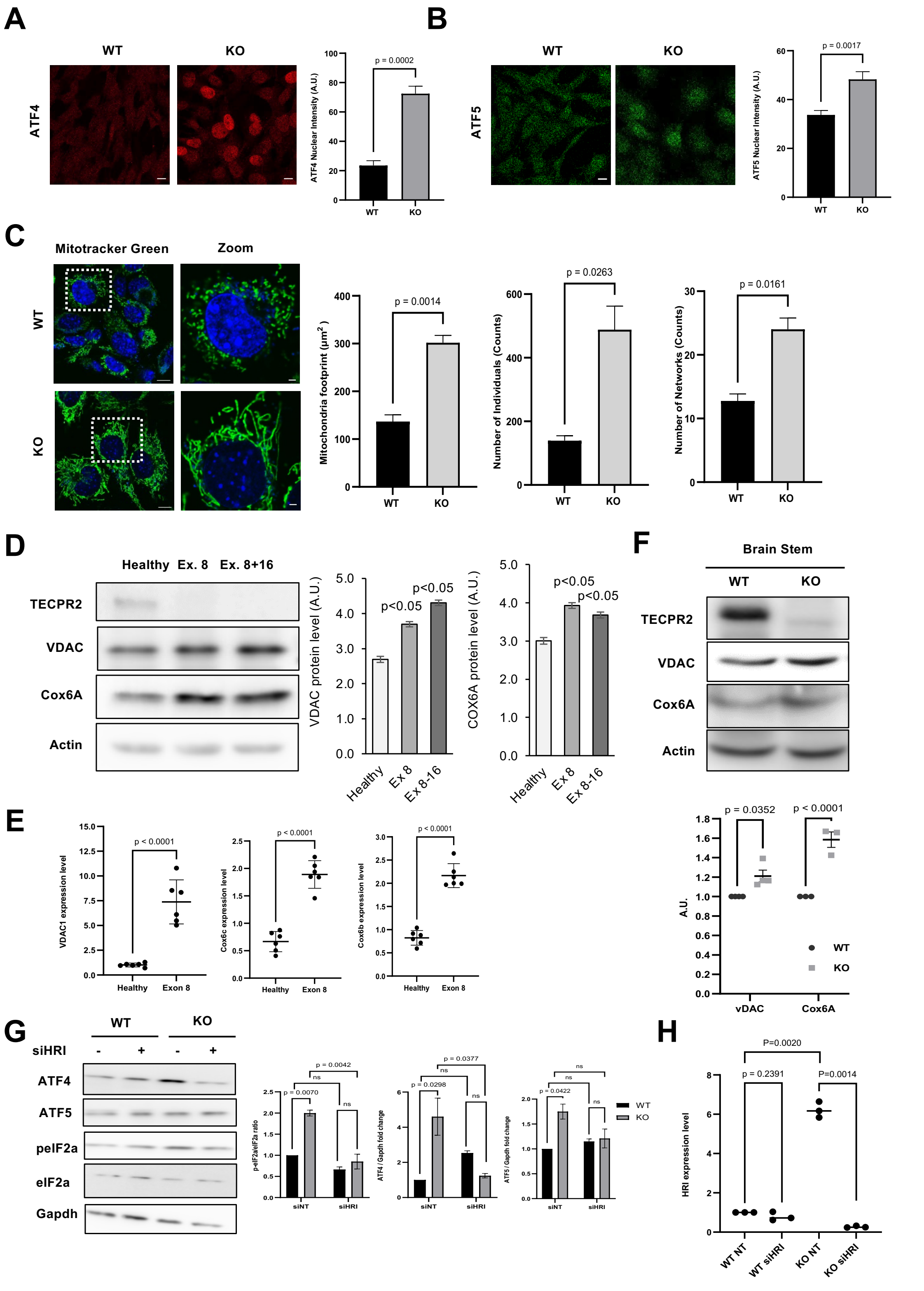


**Figure S4. TECPR2 suppresses mitochondrial stress response. A.** Confocal assessment of ATF4 in WT and TECPR2 mutant cells for ATF4 nuclear translocation in steady state conditions. Scale bar: 20 µm. **B.** Confocal visualization of ATF5 in WT and TECPR2 mutant cells for ATF5 nuclear translocation in basal conditions. Scale bar: 20 µm. **C.** Confocal microscopy live imaging of total mitochondria by Mitotracker Green staining in TECPR2 KO and WT MEF cells. Scale bar: 10 µm. Mitochondrial content parameters from three independent experiments were calculated and presented with SEM and Student's *t*-test, *p < 0.05 and **p < 0.01. **D.** Immunoblot analysis in total protein extracts of patient Ex8 and Ex 8+16 patient fibroblasts and corresponding heathy control for TECPR2, VDAC, and Cox6, and ACTA1-normalized levels of VDAC and Cox6 were calculated and presented (right panel) with the SEM of three independent experiments, *p < 0.05, determined by one-way ANOVA with posthoc Dunnett's Multiple Comparison Test. **E.** Quantitative real-time PCR analysis in healthy control and patient-derived fibroblast (Ex8+16) for mitochondrial markers VDAC1, Cox6c, and Cox6b expression level. Fold change of three independent experiments was calculated and presented with the SEM, and student's *t*-test ****p < 0.0001. **F.** Total protein extracts from the brain stem of WT and TECPR2 KO mice were analyzed by western blotting for TECPR2, VDAC, Cox 6, and Actin-normalized levels of VDAC and Cox6 were calculated and presented (right panel) with SEM of three independent experiments, *p < 0.05 for VDAC and ***p < 0.001 for Cox6A, determined by paired *t*-test analysis. **G.** Partial recovery of p-eIF2α/total eIF2α ratio and ATF4 protein level upon HRI knockdown estimated by western blot analysis. Data presented from two independent experiments. **H.** Validation of HRI knockdown by real-time PCR in TECPR2 WT and KO MEF cells performed for 72 hours using Darmophect1 reagent. Data presented from three independent experiments **p < 0.01; ns-non-significant using unpaired *t*-test analysis.

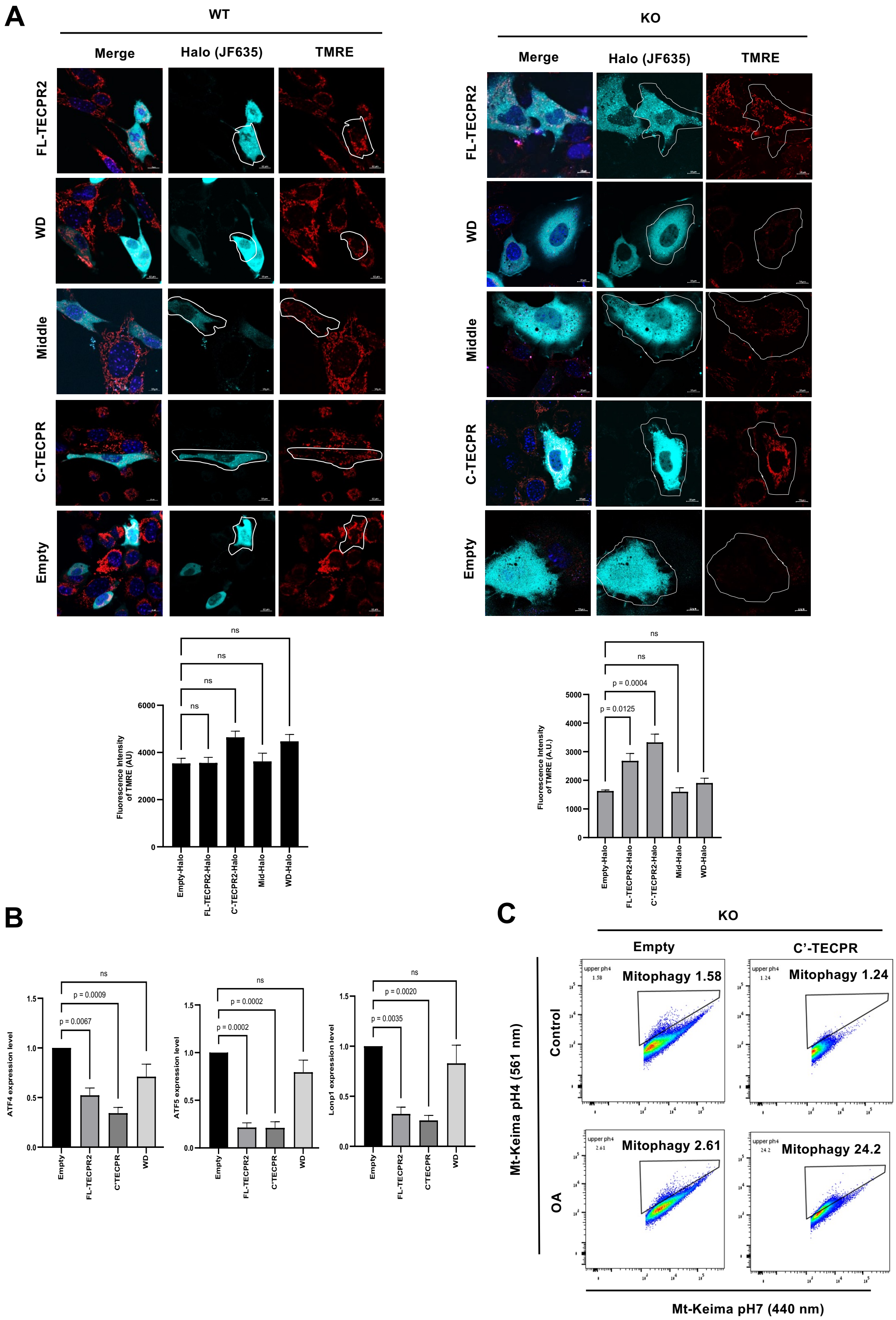


**Figure S5. The TECPR domain maintains mitochondrial homeostasis. A.** TECPR2 WT and KO MEF cells were transfected with the indicated domains for 48 hours employing JetPrime transfection reagent. Then, the cells were treated with TMRE, and confocal microscopy was used to estimate mitochondrial polarization. The mean fluorescence intensity of TMRE was calculated and presented with the SEM of three independent experiments, ***p < 0.001, **p < 0.05, and ns- non-significant, determined by one-way ANOVA with post-hoc Dunnett's multiple comparison test. **B.** Quantitative real-time PCR analysis of ATF4, ATF5, and LONP1 transcription levels in stably expressing TECPR2 domains. Fold changes were calculated and presented with the SEM of three independent experiments, ****p < 0.0001, ***p < 0.001, **p < 0.01, *p < 0.05, and ns- non-significant, determined by one-way ANOVA with post-hoc Dunnett's multiple comparison test. **C.** Density plots of FACS analysis for mitophagy flux estimation in MEF stably expressing Empty-Halo-Flag-tag cassette and C-terminal TECPR (Fig. 5D).

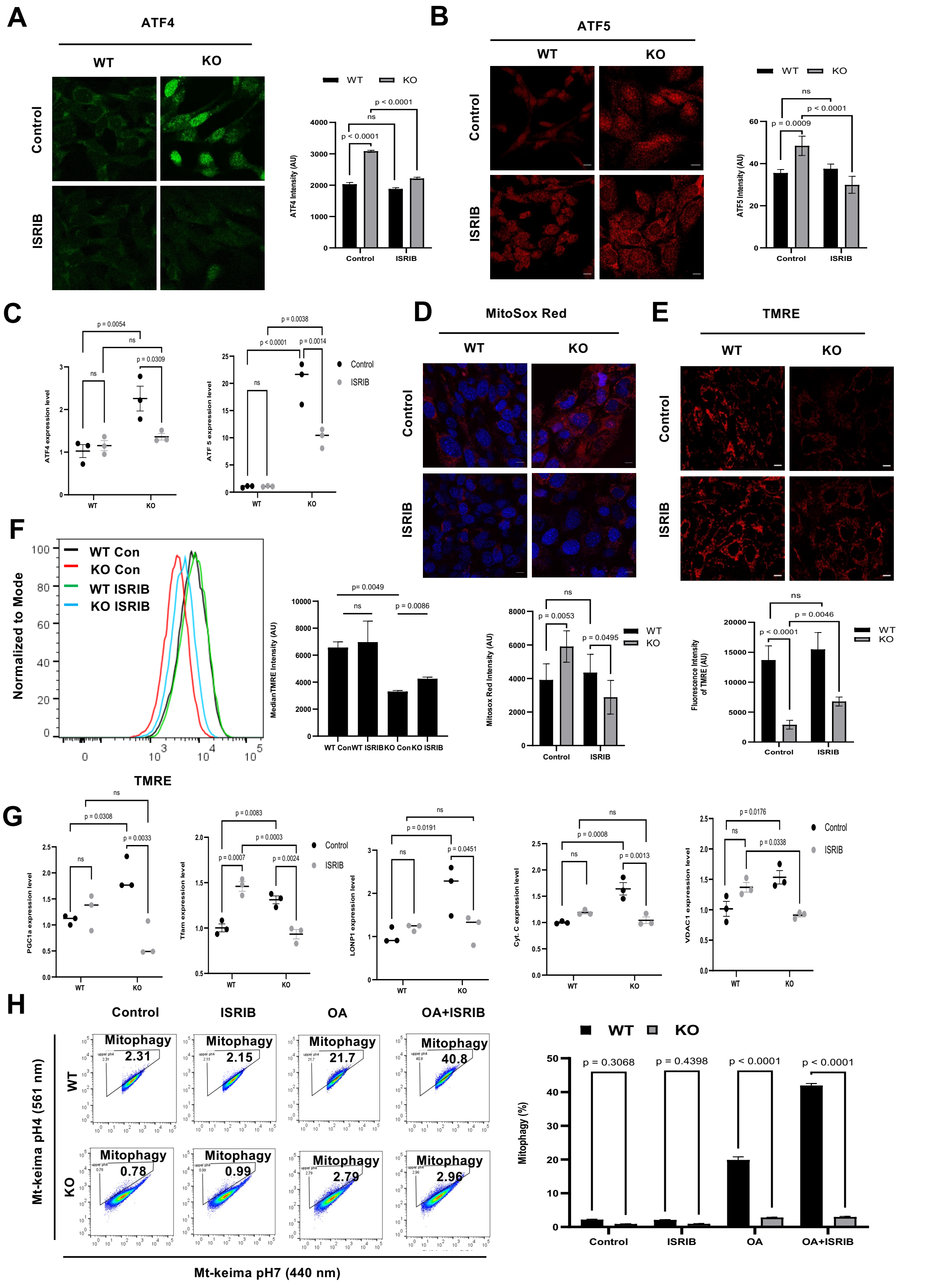


**Figure S6. ISR inhibition rescues mitochondrial function in TECPR2 mutants. A.** Confocal microscopy analysis of WT and TECPR2 KO MEF cells for ATF4 and **B.** ATF5

localizations in the presence or absence of 200nM ISRIB for 4 hours. Scale bar: 20 μm. SEM was calculated and presented (right panel) from three biological replicates. ****$p < 0.0001$ for WT and KO control, ****$p < 0.001$ between KO control and KO ISRIB treatment, respectively, using two-way ANOVA analysis with Tukey's multiple comparison test. **C.** Quantitative real-time PCR analysis in WT and TECPR2 KO MEF cells treated with 200 nM ISRIB for 4 hours for the expression level of $UPR^{mt}$ markers ATF4 and ATF5. Fold changes were calculated and presented (right panel) with the SEM of three independent experiments. Two-way ANOVA determined significance with Tukey's multiple comparison test. **D.** Live confocal microscopy analysis of mtROS in WT and TECPR2 KO MEF cells treated with 200 nM ISRIB for 4 hours. The quantitative data from three independent experiments represent mean ± SEM, **$p < 0.01$, *$p < 0.05$. **E.** Live confocal microscopy analysis of mitochondrial membrane potential upon 200 nM ISRIB for 4 hours treatment of WT and TECPR2 KO MEF cells stained with TMRE. Scale bar: 20 μm. The fluorescence intensity of TMRE was calculated and presented with the SEM of three independent experiments, using two-way ANOVA with Tukey's multiple comparison test. ****$p < 0.0001$, **$p < 0.01$, and ns for non-significant. **F.** Flow cytometry analysis of mitochondrial polarization in 200 nM ISRIB (4 hours) treated WT and TECPR2 KO MEF cells by TMRE. Median TMRE intensity was calculated from 3 independent experiments. Two-way ANOVA determined significance with Tukey's multiple comparison test. *$p < 0.05$, **$p < 0.01$, ****$p < 0.0001$, ns for non-significant. **G.** Quantitative real-time PCR analysis in WT and TECPR2 KO MEF cells treated with 200 nM ISRIB (4 hours) for PGC1, TFAM, LONP1, Cyt.C., and VDAC1 expression level. Fold change is presented from three independent experiments, representing mean ± SEM, ***$p <0.001$; **$p < 0.01$; *$p < 0.05$; ns – non-significant. **H.** Flow cytometry analysis of mitophagy in stably expressing mt-Keima WT and TECPR2 KO MEF cells pretreated with 200 nM ISRIB for 4 hours and then treated overnight with OA. Data shown from three independent biological replicates. ****$p < 0.001$ for WT OA with KO OA and ****$p < 0.001$ WT OA+ISRIB with KO OA+ISRIB treatment. Two-way ANOVA determined significance with Tukey's multiple comparison test.
